# Modeling HIV infection, treatment, rebound, and intervention in human immune organoids

**DOI:** 10.64898/2026.01.25.701611

**Authors:** Sarah A. Sackey, Jacob E. Schrum, Uma M. Mangalanathan, Rachel S. Fish, Elsa Sola, Ahmad Selehi, Ruoxi Pi, Jerome A. Zack, Jocelyn T. Kim, Dara A. Lehman, Mark M. Davis, Catherine A. Blish

**Affiliations:** Stanford Immunology Program, Stanford University School of Medicine, Stanford, CA, USA; Department of Medicine, Stanford University School of Medicine, Stanford, CA, USA; Division of Human Biology, Fred Hutch Cancer Center, Seattle, WA, USA; Institute for Immunity, Transplantation, and Infection, Stanford University School of Medicine, Stanford, CA, USA; Donor Network West, San Ramon, California, USA; Department of Microbiology, Immunology, and Molecular Genetics, University of California, Los Angeles, Los Angeles, California, USA; Department of Medicine, Division of Hematology and Oncology, University of California, Los Angeles, Los Angeles, California, USA; Department of Medicine, Division of Infectious Diseases, University of California, Los Angeles, Los Angeles, California, 90095, USA; Department of Microbiology and Immunology, Stanford University School of Medicine, Stanford, CA, USA; Chan Zuckerberg Biohub, San Francisco, CA, USA

## Abstract

Targeting the HIV-infected reservoir in lymphoid tissues (LT) will be critical to developing a cure for people living with HIV (PLWH). LT explants used to study HIV infection enable the evaluation of human-specific disease progression and treatment response; however, their short lifespan makes it challenging to assess long-term treatment interventions. We therefore established an immune organoid model of HIV infection using human tonsil or spleen cells, demonstrating productive HIV infection and viral integration into CD4^+^ T cells. Treatment with a protease inhibitor fully suppressed ongoing viral production, with virologic rebound occurring within days of treatment interruption. The transfer of healthy allogeneic NK cells to target the reservoir upon treatment interruption reduced the number of infected cells, intact viral genomes, and production of *de novo* infectious viral particles. Adoption of this immune organoid platform will accelerate the evaluation of cure-based strategies to eliminate the HIV reservoir in tissues for PLWH.

## Introduction

Currently, 40 million people are living with human immunodeficiency virus (HIV), with over a million new cases annually ^1^. Post-infection integration of viral DNA (vDNA) establishes a reservoir of infected cells in the blood, brain, and lymphoid tissue (LT) compartments ^2–5^. The highest burden of infected cells are found in LTs, including the lymph node, gut-associated lymphoid tissue, and spleen ^5,6^. For people living with HIV (PLWH), treatment in the form of antiretroviral therapy (ART) inhibits viral replication and suppresses the presence of viral RNA (vRNA) in blood below the limit of detection by clinical assays; however, ART does not eliminate the reservoir of infected cells in blood or tissues ^5,6^. ART is generally lifelong, as viral rebound typically occurs within 5-21 days following analytical treatment interruption (ATI) ^7–10^. This rebound is driven by the underlying viral transcription in the reservoir of infected cells in tissues, even in the presence of ART ^3–6,11,12^. The majority of reservoir and rebound studies focus on blood as this is an easier specimen to collect from study participants; however, a cure will require targeted elimination of the tissue reservoir.

Evaluation of potential HIV cure strategies use *in vitro* models such as HIV infected cell lines, primary CD4^+^ T cells isolated from blood, and *in vivo* animal models. Each of these models presents advantages and limitations. The *in vitro* systems provide an understanding of infection and reservoir in the blood ^13,14^, HIV permissive cells ^15–17^, evaluation of ART ^18–20^, and cytolytic function against infected cells ^21^, but they fail to capture the tissue-specific characteristics of infection. This presents the primary advantage of *in vivo* models: mice and non-human primates (NHPs) allow the evaluation of infection at the whole-organism level, capturing infection in tissues. However, these animals do not inherently support HIV infection. Mice must be humanized by providing human immune cells to make them amenable for the study of HIV replication, ART and latency ^22^, and cure strategies ^23^. However, the humanization process is expensive, labor-intensive, and imperfect. Humanized mice often have deficiencies in humoral development and in reconstitution of innate immune cells, limiting the study of viral spread in the myeloid population ^24–26^, and the mice can develop graft-versus-host disease ^27^. Non-human primates (NHPs) provide an alternative model, with selected strains and species exhibiting disease progression similar to that observed in humans ^28,29^. In particular, macaques show a similar disease progression as humans when infected with simian immunodeficiency virus (SIV)^29^. However, evolutionary differences between SIV and HIV limit the use of HIV-1 in NHPs. Some of these limitations including the use of some antiretrovirals (ARVs) and the development of antibodies have been overcome by developing simian-human immunodeficiency virus (SHIVs); however, translation of results from NHP studies to humans have not always been successful ^29–33^. NHP use is also expensive, tightly regulated, and can raise ethical concerns, limiting the size and scale of NHP studies ^33,34^. Thus, there remains a need for a cost-effective, high-throughput, pathobiologically relevant system with high fidelity to human infection in tissues to drive HIV cure research.

To better model the complex cellular composition and cellular interactions in human secondary lymphoid tissues, human lymphoid histocultures (HLH), first established in 1995 ^35,36^, and human lymphoid aggregate cultures (HLACs), introduced in 2001, addressed this need ^37^. These cultures, derived from tonsil and adenoid tissues, provided valuable insights into infection kinetics in tissues using primary HIV isolates and lab-adapted HIV strains; however, these tissues are cultured for less than 2 weeks due to poor viability ^37,38^. These *ex vivo* cultures provide insight into the permissive nature of memory cells ^39^, germinal center T follicular helper cells ^40^, determinants of abortive and productive infection in various cell types ^41^, viral accessory protein function ^42^, and mechanisms of cell death and inflammation during infection ^17,38,43,44^. Although the HLH and HLAC systems provided these key insights to viral infection in tissues, they have yet to mimic other key features of disease progression, such as virologic rebound. Herein, we extend that research with tonsil and spleen immune organoids, which were initially developed to evaluate vaccine responses ^45^. Unlike the HLACs, these organoids are cultured in transwells in the presence of BAFF, a key cytokine essential for maintaining culture viability, antibody production, and spontaneously self-assemble within 3 days into three-dimensional (3-D) lymphoid structures, including germinal centers ^45^. They approximate normal lymphatic organs, as evidenced by the fact that they can respond to vaccines for influenza through the proliferation of antigen-specific B and T cells, somatic mutation, affinity maturation, and class switching ^45^. Using a replication-competent early infection CCR5-tropic subtype A strain of HIV, Q23 ^46^, we benchmarked the system’s ability to recapitulate pathobiological features observed in PLWH, including the productive infection of CD4^+^ T cells, identification of p24^+^ myeloid cells, inhibition of viral replication with ART, and virologic rebound post ATI. Most importantly, we tested the effects of a latency-reversing agent (LRA) and a natural killer (NK) cell-based cure strategy. We show that NK cell intervention reduces viral output and the number of infected cells, results parallel to those observed in a humanized mouse model ^23^. Overall, we demonstrate that immune organoids mimic *in vivo* infection, rebound, and intervention and can serve as a platform to test other cure based approaches for PLWH.

## Results

### Establishment of HIV infection in tonsil and splenic organoids

To model HIV infection in the 3-D immune organoids, we assessed the ability of organoids derived from CD45^+^ tonsillar or splenic cells to support HIV infection across a multiplicity of infection (MOI) range from 0.001-10, using the HIV strain Q23. We tested infection before (on day 0) and after (on day 3) organoid aggregation (Table, 1, Fig. 1 and Supplementary Fig. 1). Organoids aggregated normally when infected on day 0 and evaluated 3 days post-infection (dpi) (Fig. 1b). We quantified infection by measuring the frequency of Gag p24^+^ cells among CD3^+^CD8^-^ T cells to account for CD4 downregulation upon infection (Supplementary Fig. 1a). In the tonsil organoids infected on day 0, we observed significant levels of infection at MOI 0.1, with a mean frequency of 2.8% (range, 0.25-8.7%) of p24^+^ CD3^+^CD8^-^ T cells. The frequency of infected cells did not increase significantly with higher MOI (Fig. 1c), suggesting that an MOI of 0.1 was sufficient to infect most of the HIV-permissive cells. In the splenic organoids we observed an average 1.7% (range, 0.8-2.9%) p24^+^ CD3^+^CD8^-^ T cells at an MOI of 0.1 and subsequent increase in infection at mean of 3.4% (range 3.1-4.1%) at an MOI of 1, but infection levels did not increase further at an MOI of 10 (Fig. 1d). Similar results were observed when infection was performed on day 3, following organoid aggregation, with corresponding analysis on day 6 (Supplementary Fig. 1b-d). Interestingly, the two tonsil organoids derived from pediatric tissues consistently exhibited higher infection rates than adult-derived organoids (Fig. 1c and Supplementary Fig. 1c). Based on these results, an MOI of 0.1 was used in subsequent experiments.

**Table 1:** Donor demographics of all the tonsil spleen derived tissues used in this work.

|  | Tonsil<br>(N=16) | Spleen<br>(N=17) |
| --- | --- | --- |
| **Age (years)** |  |  |
| Mean (SD) | 22.2 (12.4) | 47.6 (11.3) |
| Median [Min, Max] | 22.0 [7.00, 49.0] | 51.0 [27.0, 62.0] |
| **Sex** |  |  |
| Male | 10 (62.5%) | 10 (58.8%) |
| Female | 6 (37.5%) | 7 (41.2%) |
| **Race/Ethnicity** |  |  |
| Asian | 2 (12.5%) | 4 (23.5%) |
| Black or African American | 0 (0%) | 2 (11.8%) |
| Hispanic/Latino | 7 (43.8%) | 2 (11.8%) |
| White - Non-Hispanic/Latino | 5 (31.3%) | 0 (0%) |
| White | 0 (0%) | 9 (52.9%) |
| N/A | 2 (12.5%) | 0 (0%) |
| **Status** |  |  |
| Deceased | 0 (0%) | 17 (100%) |
| N/A | 16 (100%) | 0 (0%) |

**Fig. 1:**
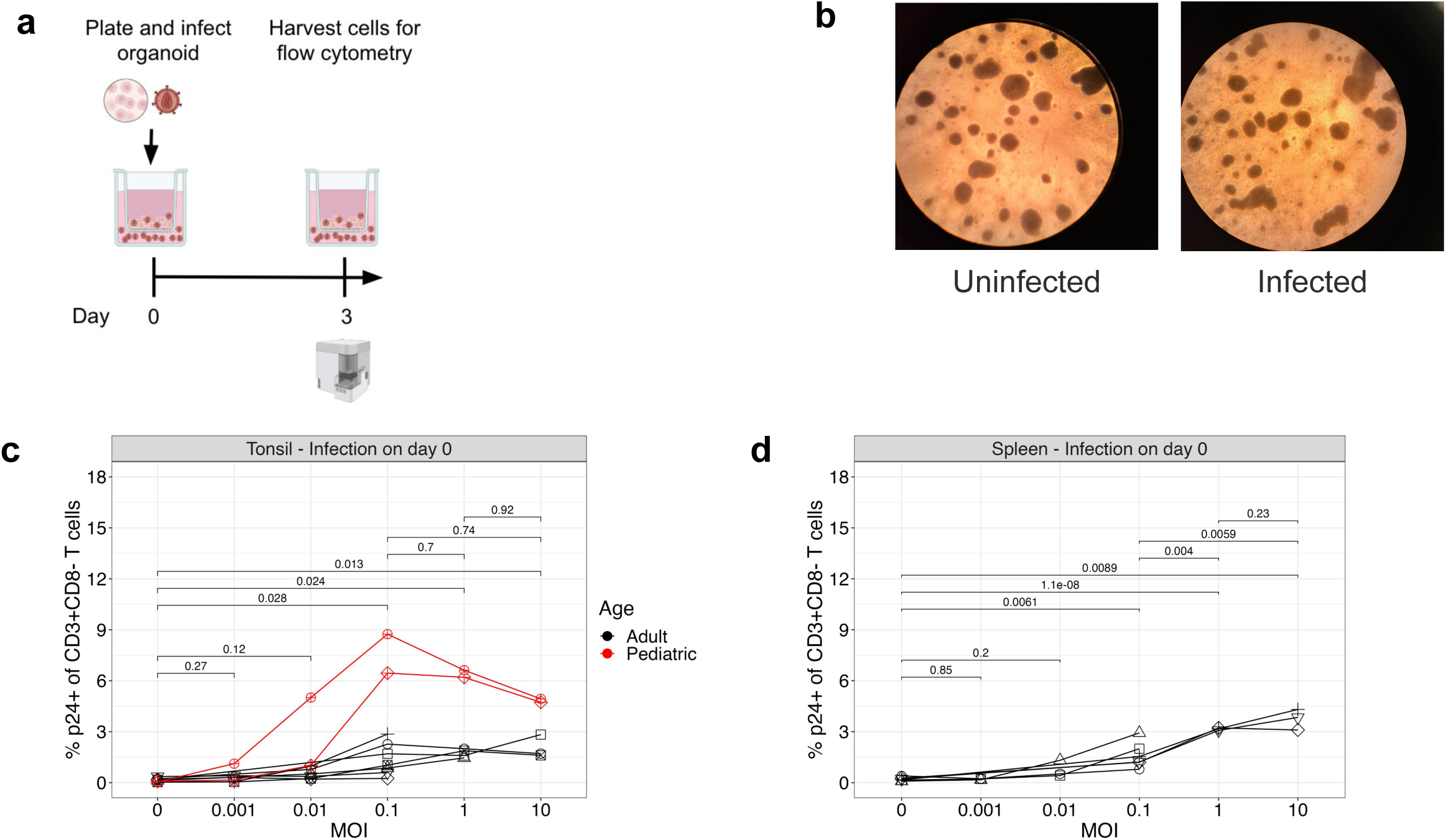
Limited population of tissue cells is permissive to HIV infection. **a**, Schematic of infection timeline. Organoids were infected on day 0 then harvested for flow cytometry 3 dpi. **b**, Images 3dpi of uninfected and infected tonsil organoid at 4X magnification. **c-d**, Titration of MOI in **c**, tonsil and, **d**, splenic organoids, measuring frequency of HIV Gag p24+ CD3+CD8- T cells 3 dpi. Tonsil organoids were derived from 9 donors, 7 adult (black) and 2 pediatric (red). Splenic organoids were derived from 6 adult donors. Each symbol represents a donor. Statistical analysis in were performed with unpaired t test.

Importantly, infection occurred in the absence of exogenous stimulation. This is in stark contrast to infection of T cells isolated from PBMCs, which require stimulation with PHA, anti-CD3 and anti-CD28 to render them permissive to infection. These activated CD4^+^ T cells from blood required an MOI of 1-5 to achieve a similar range of infection as observed in the organoids at a MOI of 0.1 (Supplementary Fig. 1e). This suggests that tissue-derived immune organoids are intrinsically permissive to infection.

### Antiretroviral treatment of tonsil and splenic organoids

To model treatment while ensuring successful first-round infection and integration, we targeted the end of the viral life cycle with tipranavir (TPV), a protease inhibitor (PI). Previous studies determined the 90% inhibitory concentration (IC90) of PNU-140690, later named tipranavir, 22against clinical isolates to be between 0.1 and 0.8 μM when tested in isolated CD4^+^ T cells ^18,19^. TPV was added at time of infection and titrated across a concentration range from 0.01 to 10 μM and evaluated 3 dpi. Treatment with the PI did not affect the rate of infection (Supplementary Fig. 2a). Suppression of *de novo* infectious particles (IP) was achieved at at an IC90 of 10 μM (Supplementary Fig. 2b). This concentration was used in subsequent experiments.

To determine if this organoid system is able to model the reduced inflammation that occurs *in vivo* during ART, we measured CXCL10, also known as interferon gamma-induced protein (IP-10) because CXCL10 levels have been associated with treatment response and disease progression ^47–49^. HIV infection trended towards increased CXCL10 expression at 3 dpi, which was reduced with TPV (Fig. 2A).

**Fig. 2:**
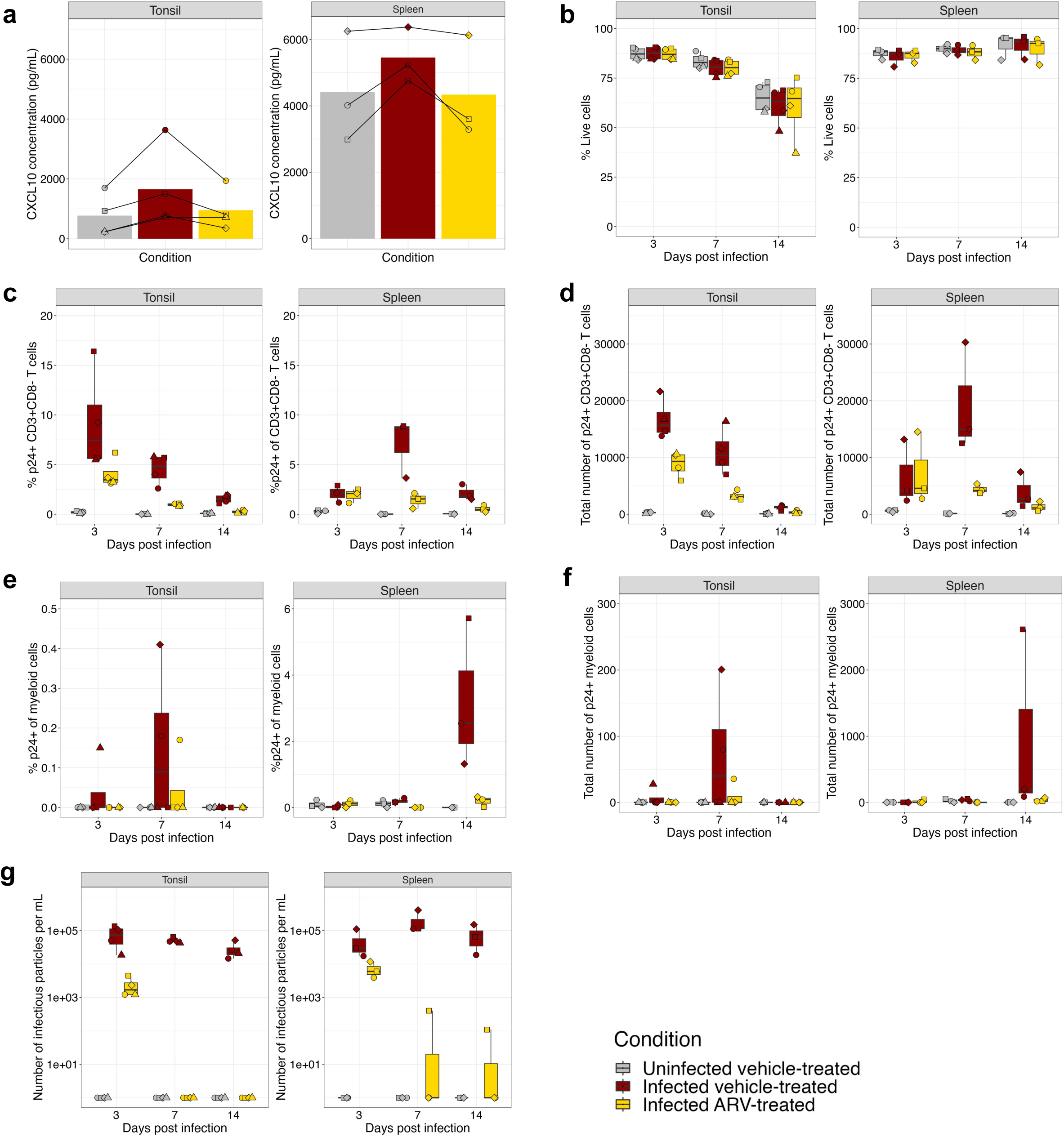
ART suppressed inflammation, infection, and production of *de novo* viral particles. 14 day time course infection of tonsil and splenic organoids. **a**, CXCL10 concentration of organoid supernatants at 3dpi. **b-f**, 14 day time course infection of tonsil and splenic organoids. Frequency of, **b**, Live, **c**, p24+ CD3+CD8-, and, **e**, p24+ myeloid cells. Quantification of the number of, **d**, p24+ CD3+CD8- and, **f**, p24+ myeloid cells. **g**, Quantification of the number of infectious viral particles in the supernatant of the organoids on the TZM-bl cell line. Tonsil organoids were derived from 4 pediatric donors, splenic organoids were derived from 3 adult donors. Each symbol represents a donor and is the mean of technical duplicates.

Next, we investigated cellular composition and production of IP post infection over a 14-day time course (Fig. 2 and Supplementary Fig. 3). In the tonsil organoid, viability decreased from a mean of 87% at 3 dpi to 60% at 14 dpi (Fig. 2b). In contrast, the splenic organoids maintained excellent viability over 14 days in culture, consistently above 85% across infection and treatment conditions (Fig. 2b). The overall frequency of major cell subsets CD19^+^ B cells, CD3^+^CD8^-^ T cells, CD3^+^CD8^+^ T cells, NK cells, and myeloid cells did not change substantially over time in uninfected cultures (Supplementary Fig. 3 and Supplementary Fig. 4a). There was notable depletion of CD3^+^CD8^-^ T cells in infected, vehicle-treated organoids, particularly at days 7 and 14 (Supplementary Fig. 4a).

### Infection kinetics differ between tonsil and splenic organoids

Quantification of infection demonstrated that ART reduced the frequency and number of p24^+^ CD3^+^CD8^-^ T cells to levels to near undetectable levels by 14 dpi in both tonsillar and splenic organoids (Fig. 2c,d). Interestingly, we observed differences in infection and treatment kinetics between the two tissue organoids. In the tonsils, peak infection was observed 3 dpi, with a mean of 9.2% or 1.7*10^4^ p24^+^ CD3^+^CD8^-^ T cells, while in the splenic organoids, peak infection was observed 7 dpi, with a mean of 7.1% or 1.9*10^4^ p24^+^ CD3^+^CD8^-^ T cells (Fig. 2c). In the ART condition, peak infection was observed at 3 dpi in both tissues, mean infection rate of 4% or 8.8*10^3^ in tonsil and 1.9% or 7.3*10^3^ p24^+^ CD3^+^CD8^-^ T cells in the spleen (Fig. 2c,d).

### The p24^+^ T cell population is primarily composed of CM, EM, and T_fh_

To better understand which CD3^+^CD8^-^ T cells types were infected, we evaluated the frequency of major CD3^+^CD8^-^ T cells subsets, including germinal center (GC) (CXCR5^hi^PD1^hi^), and T follicular helper cells (T_fh_) (CXCR5^+^PD1^+^), central memory (CM) (CD45RA^-^CCR7^+^), effector memory (EM) (CD45RA^-^CCR7^-^), naive (CD45RA^+^CCR7^+^), effector memory cells re-expressing CD45RA (TEMRA) (CD45RA^+^CCR7^-^) among total CD3^+^CD8^-^ T cells in the uninfected or the infected cultures stratifying by p24^-^ (bystander) and p24^+^ (infected) (Supplementary Fig. 3 and Supplementary Fig. 4b). The phenotype of cells within the p24^+^ fraction of both vehicle-treated and ARV-treated cultures was almost entirely composed of CM, EM, and TFH cells (Fig. S3 and S4B). Over the 14-day experiment, in tonsil-derived organoids, the mean frequencies of major CD3^+^CD8^-^ T cell subsets from the p24^+^ fraction were 42.8-52.8% EM, 33.9-34.0% CM, and 11.9-17.5% T_fh_ cells. In the spleens, the frequencies of these subsets were 16.3-51.6% EM, 24.6-67.8% CM, and 4.1-8.6% T_fh_ cells, with notable contribution from TEMRA subset 5.8-16.9%. Similar frequencies of EM, CM, and T_fh_ were observed in the p24^+^ fraction of the infected ARV-treated conditions in both tissues (Supplementary Fig. 4b). Together, these findings demonstrate that memory and T_fh_ cells are the primary cell types initially infected and are maintained over time. This is consistent with previous findings that these cell types constitute the majority of the reservoir ^11,15,16,40^.

### CD8^+^ T cells and NK cells are activated and proliferative in response to infection

We next examined activation of effector CD8^+^ T cells and NK cells post-infection. Infection increased the frequency of HLA-DR^+^CD8^+^ T cells, which was partially reduced by ARV (Supplementary Fig. 5a); no such trend was observed in NK cells (Supplementary Fig. 5b). The frequencies of Ki-67^+^CD8^+^ T cells and NK cells were more variable, with a trend toward an increase at day 7 post-infection in the tonsils (Supplementary Fig. 5c,d). Overall, these data suggest that HIV infection activated effector cells in some donors, and this activation was reduced by ART.

### Identification of p24^+^ myeloid cells in immune organoids

We observed a low frequency of p24^+^ cells in the myeloid fraction at 14 dpi in the spleen, with an average of 3.2% p24^+^ myeloid cells, and an even smaller fraction, 0.15%, in the tonsils, although this was inconsistently detected (Fig. 2e and Supplementary Fig. 6a). There was wide variability in the absolute number of total myeloid cells and p24^+^ myeloid cells in tonsil and spleen from a few hundred to several thousand (Fig. 2f). Backgating confirmed that the p24^+^ myeloid population clustered at higher forward- and side-scatter, characteristic of myeloid cells (Supplementary Fig. 6a).

### Antiretroviral treatment suppresses the production of infectious particles

Finally, we quantified *de novo* virion production in the cultures using a TZM-bl assay. The infected, vehicle-treated immune organoids produced 2.8*10^4^-2.1*10^5^ IP/mL between 3-14 dpi. ARV added at time of infection reduced the IP to an average of 2.3*10^3^ in tonsils and 7.2*10^3^ in spleens at 3 dpi and completely suppressed IP production by 7 dpi in all of the tonsil donors and 2 out of 3 of the splenic donors (Fig. 2g). Although not fully suppressed, this one splenic organoid produced ∼600-fold less virus than its infected, vehicle-treated counterpart by day 14 (Fig. 2g). Thus, we achieved total suppression of infectious particles one week post-treatment in the majority of infected immune organoids.

### Visualization of HIV-infected cells via RNAscope-FISH

To directly visualize HIV-infected cells in the organoids, we used RNAscope fluorescence in situ hybridization (RNAscope-FISH) for vDNA and/or vRNA ^6,50,51^. RNAscope-FISH was performed on organoids grown using either traditional transwell inserts (Fig. 3a), or in chambered slides (Fig. 3b), which were used because sectioning traditional transwell-grown organoids led to shear stress and loss of architecture on many sections. In the traditional organoids, we stained only for vDNA and observed several cells with several copies of HIV DNA, suggestive of integrated provirus and/or unintegrated form of HIV DNA generated during replication by 3 dpi (Fig. 3a). For the organoids grown in chambered slides, we observed co-localization of CD4 and vDNA (Fig. 3b, white arrow) as well as co-localization of vRNA and vDNA (Fig. 3b, red arrow). Viral packaging occurs at the cell membrane ^52^; thus, the detection of viral Env RNA at the cell membrane indicates active vRNA transcription and viral packaging at the membrane. Organoids grown in chambered slides produced similar quantities of IP as those grown in traditional transwells (Fig. 3c and Fig. 2g) and average of 10^4^-10^5^ IP/mL, establishing this as a viable alternative method for growing organoids for imaging.

**Fig. 3:**
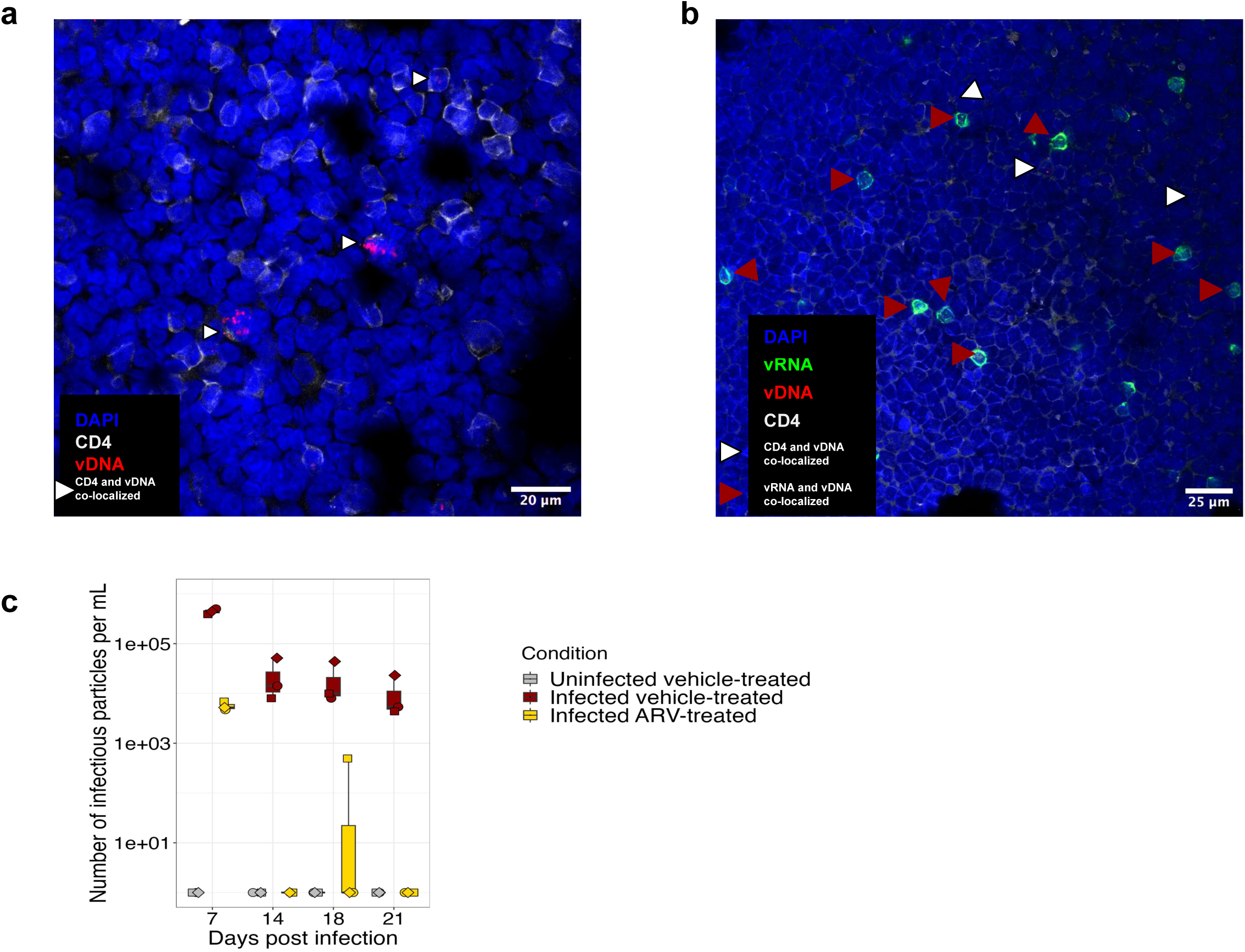
Organoids are stable for microscopy by sectioning or grown in chambered slide. Representative confocal microscopy images of RNAscope-FISH of splenic organoids grown in, **a**, transwell, **b**, chambered slide 3-4 dpi. **a**, Anti-CD4 antibody in gray along with predesign HIV DNA sense probes in red, 3 dpi (magnification 63X, oil). **b**, Anti-CD4 antibody in gray along with custom probes against HIV-Q23, sense probes (targeting vDNA) in red, and antisense probes (targeting vRNA) in green, 4 dpi. **c**, Quantification of the number of infectious viral particles in the supernatant of the organoids grown in a chambered slide at 7, 14, 18, 21 dpi. Each symbol represents a donor and is the mean of technical duplicates.

### Rebound occurs within one week of ATI

We next explored whether our organoids could model the kinetics of viral rebound following ATI. Given the plateau in infection with increasing MOI (Fig. 1), we understood most of the HIV-permissive cells were likely infected in the first round of infection; thus, we reasoned that we would need to mimic the migration of uninfected target cells to the site of infection by adding fresh donor-matched uninfected splenocytes. Therefore, we infected and TPV-treated the cultures for 7 days to suppress viral replication. On day 7 (Day 0 ATI), we removed 50% of the cells for phenotyping and replaced them with 3*10^6^ fresh donor-matched uninfected splenocytes. Concurrently, we replaced the media to remove TPV and followed infection for another 7 days (Fig. 4a). In infected, vehicle-treated conditions, we observed inter-individual variation in cell frequencies, with depletion of CD3^+^CD8^-^ T cells on day 0 and 7 post ATI (Fig. 4b). The ATI condition also saw slight depression in CD3^+^CD8^-^ T cells (Fig. 4b). At 7 days post ATI, there was a slight increase in the frequency of p24^+^ CD3^+^CD8^-^ T cells in the ATI condition, compared to the ART condition, suggestive of rebound (Fig. 4c). We detected no IP in the supernatant on ATI day 0, followed by an average of ∼3.5*10^3^ and ∼1.5*10^4^ IP/mL in the supernatant at 3 and 7 days post ATI respectively. In the cultures that maintained ART, we did not detect any IP (Fig. 4d). The steady increase in infectious *de novo* virions in supernatant over time demonstrates successful viral rebound.

**Fig. 4:**
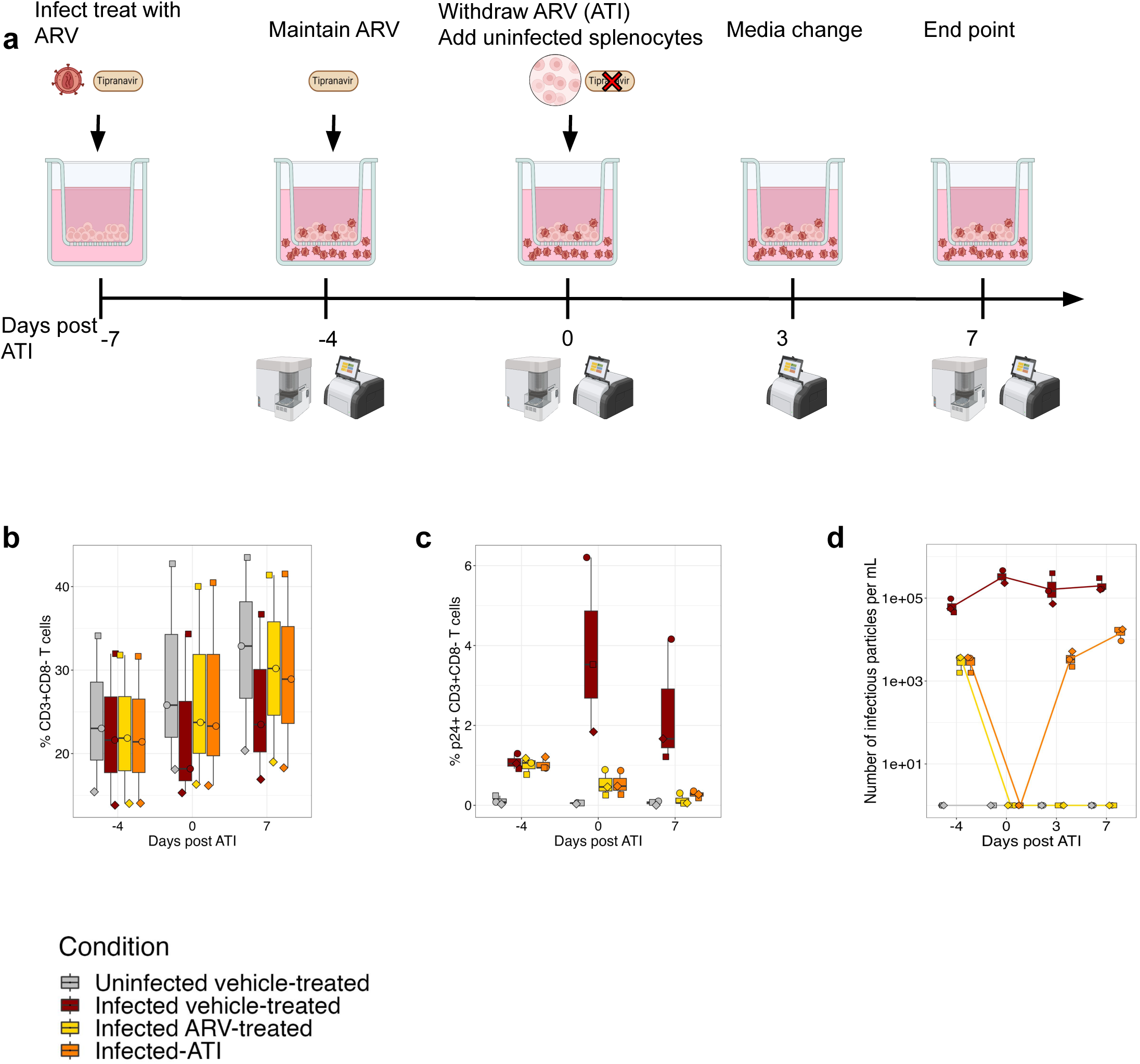
Addition of uninfected splenocytes allows for robust rebound. **a**, Schematic of the culture timeline to induce rebound measured by days post-analytical treatment interruption (ATI). Frequency of, **b**, CD3+CD8- cells and, **c**, p24+ CD3+CD8- cells. **d**, Quantification of the number of infectious viral particles in the supernatant of the organoids post ATI. Splenic organoids were derived from 3 adult donors. Each symbol represents a donor and is the mean of technical duplicates.

### Transfer of allogeneic NK cells reduces viral reservoir and production of infectious particles

Finally, we used the organoid system as a platform to test an NK cell-based ‘kick and kill’ strategy for HIV cure ^53^, similar to one previously performed in a humanized mouse model ^23^. This prior study demonstrated that adoptive transfer of allogeneic NK cells at ATI delayed viral rebound and reduced reservoir size ^23^. Thus, we usedallogeneic NK cells as cytotoxic effector cells and IL-15 as a support for NK cell survival and cytotoxicity and as a the LRA via inducing of cytokine-mediated proviral transcription ^54^. This experiment consisted of six conditions: uninfected, infected vehicle-treated, infected ARV-treated, infected ATI (ATI), infected ATI with IL-15 (IL-15 alone), and infected ATI with IL-15 and NK cells (NK condition) (Fig. 5a). The experimental layout was identical to the rebound experiment, with the addition of 5*10^5^ NK cells as appropriate. In conditions with IL-15 and/or NK cells, IL-15 was supplemented into the media, and NK cells were added directly into the transwell on ATI days 0 and 7 (Fig. 5a). The organoids were followed for 21 days (7 days before and 14 days post-ATI).

**Fig. 5:**
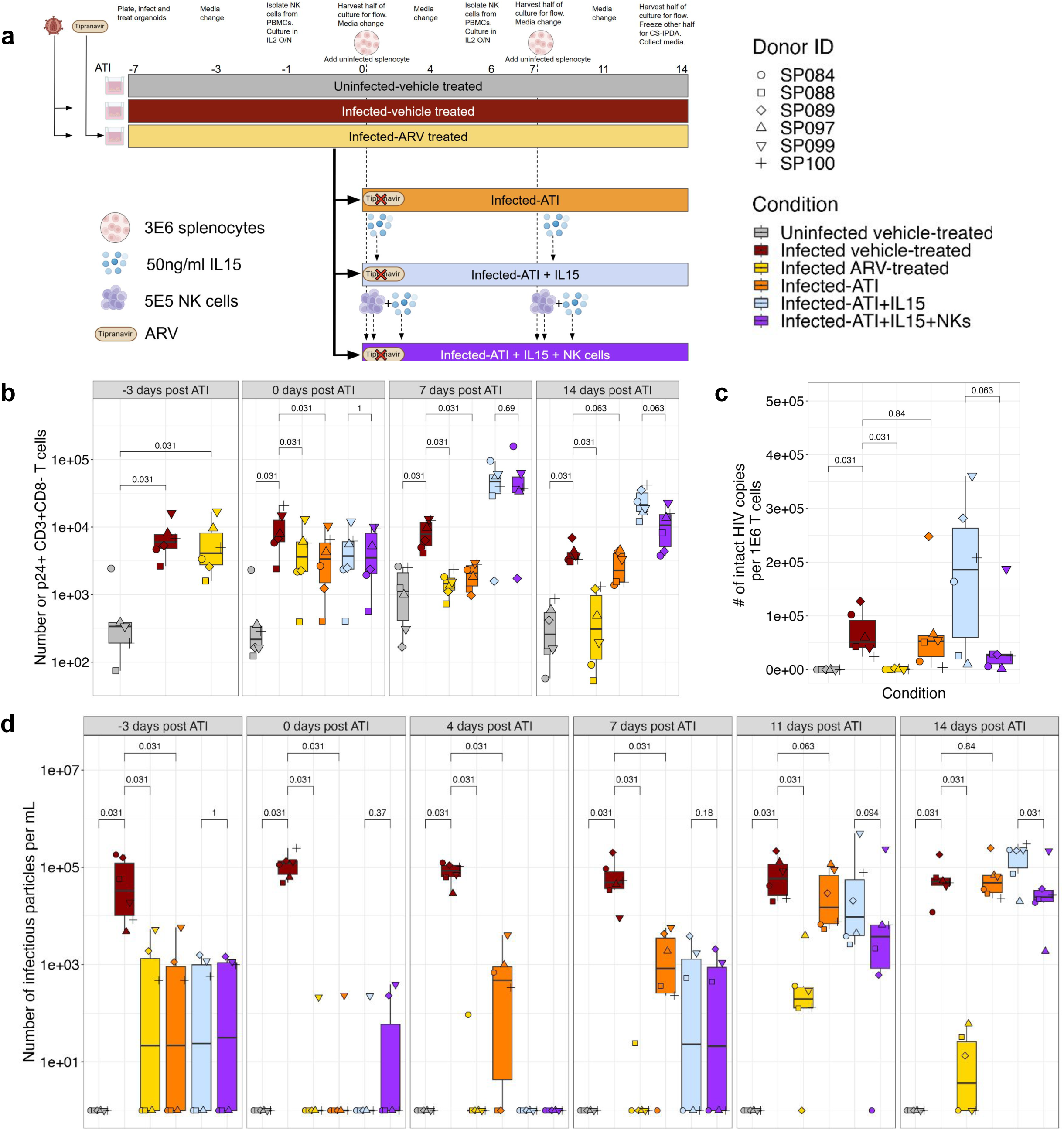
NK cell treatment reduces number of intact HIV copies and suppresses viral replication. ATI of HIV-infected splenic organoids followed by 14 days of allogeneic NK intervention. **a**, Schematic of timeline of allogeneic NK cell transfer by days post-ATI. **b**, Quantification of the total number of p24+ CD3+CD8- T cells. **c**, Quantification of intact provirus by CS-IPDA. **d**, Quantification of the number of infectious viral particles in the supernatant of the organoids by days post-ATI, with and without NK cell intervention. Splenic organoids were derived from 6 adult donors. Each symbol represents a donor and is the mean of technical duplicates. Statistical analysis in were performed with wilcoxon signed rank test.

Treatment with IL-15 did not significantly affect culture viability (Supplementary Fig. 7a). Regardless of treatment condition, infection significantly reduced the frequency of CD3^+^CD8^-^ T cells compared to the uninfected condition (Supplementary Fig. 7b). By 14 days post ATI, treatment with IL-15 enhanced activation and significantly increased the proliferation of CD3^+^CD8^-^ T cells, CD3^+^CD8^+^ T cells, and NK cells (Supplementary Fig. 7c-h).

At ATI day 0, the average frequency of p24^+^ CD3^+^CD8^-^ T cells was 1.9% (range, 0.4- 2.9%) in the infected, vehicle-treated condition; all ARV-receiving conditions had an average of less than 1% (range, 0.06-2.4%) p24^+^ CD3^+^CD8^-^ T cells (Supplementary Fig. 8a). This frequency correlated with the average number of infected cells across conditions: 9*10^3^ cells in the infected vehicle-treated condition, and 4.9*10^3^, 4.2*10^3^, 4.8*10^3^, and 4.9*10^3^ in the ART, ATI, IL-15 alone, NK condition, respectively. The consistent number of p24^+^ cells across all ARV-treated conditions at ATI day 0 demonstrates the high reproducibility of infection levels in organoids treated identically up to that point (Fig. 5b).

We observed a progressive increase in the number and frequency of p24^+^ CD3^+^CD8^-^ T cells in the ATI alone condition 7 and 14 days post-ATI. By ATI day 14, the mean frequency of infected cells in the ATI alone group was identical to that in the infected, vehicle-treated condition, at 1.5% p24^+^ CD3^+^CD8^-^ T cells. IL-15 treatment increased the frequency and number of p24^+^ CD3^+^CD8^-^ T cells compared with non-IL-15-treated conditions at 7 and 14 days post-ATI. This was to be expected given the strong stimulatory effects of IL-15; however, we hypothesized that this stimulation would render the infected cells more visible to NK cells, thereby facilitating suppression of infection. By 14 days post-ATI, treatment with NK cells reduced the average number of infected cells to 1.1*10^4^ (range, 3.7*10^3^-2.3*10^4^) from 2.4*10^4^(range, 1.2*10^4^-4.2*10^4^) in the IL-15 alone condition, representing a 54% reduction (Fig. 5b).

To directly evaluate reservoir size at the final time point, we used the cross-subtype intact proviral DNA assay (CS-IPDA) to quantify both total and intact viral genomes ^55^. Intact viral genomes are an estimate of the replication-competent proviruses, and total viral genomes account for both intact and defective proviruses. In the ARV condition, the intact reservoir size was significantly smaller than all other infected conditions, with an average of 7.9*10^3^ total HIV copies and 9.2*10^2^ intact viral genomes (Supplementary Fig. 8b and Fig. 5c). We detected an average of 6.6*10^4^ and 7.3*10^4^ intact HIV copies in the infected vehicle-treated and the ATI conditions, respectively. Treatment with allogeneic NK cells reduced the average number of both total and intact HIV copies compared with the IL-15 alone condition. The average total number of HIV copies reduced from 3.4*10^5^ in the IL-15 alone condition to 1.3*10^5^ in the NK cell-treated condition (Supplementary Fig. 8b). Accordingly, the mean intact provirus decreased from 1.7*10^5^ in the IL-15 alone condition to 4.6*10^4^ in the NK condition (Fig. 5c). NK cell treatment thus reduced the number of intact viral genomes by 73% compared with IL-15 alone. Reduction in both total and intact HIV copies was observed in all but one donor, suggesting NK cell-mediated pressure eliminates cells harboring intact vDNA.

In evaluating virus production, there was heterogeneity among donors: one donor failed to achieve complete viral suppression, and another showed a blip in the NK added condition on ATI day 0. In these two donors, viral production was ∼600-fold lower than in the infected, vehicle-conditioned group. By ATI day 4, we detected IP in the supernatant for four of the six donors in the ATI condition, but there was no detectable virus in the IL-15-treated conditions. From ATI day 7 onward, we detected virus in the supernatant for all ATI conditions. On ATI day 14, addition of NK cells reduced the the mean IP/mL from 1.8*10^5^ to 5.4*10^4^ compared to the IL-15 alone, a 70% reduction (Fig. 5d). This reduction was consistent with our hypothesis that NK cells can target HIV-infected cells and limit viral rebound. Overall, NK cell intervention reduced IP production relative to the infected vehicle-treated group in four of the six donors (SP088, SP089, SP097, SP100) (Supplementary Fig. 8c).

## Discussion

Targeted elimination of HIV-infected cells in lymphoid tissues, such as lymph nodes, GALT, the spleen, and tonsils, is essential to developing a cure for PLWH. Here, we describe an *in vitro* immune organoid system that is highly amenable to manipulation and experimentation and models key features of HIV infection, suppression, rebound, and treatment, enabling the assessment of HIV cure strategies. HIV infection of these immune organoids occurs primarily in EM, CM, and T_fh_ CD4^+^ T cells in the absence of exogenous stimulation, reflecting the cell types infected in PWLH. Other key features of natural HIV infection are replicated, including the induction of an inflammatory response and loss CD4^+^ T cells, which are partially reversed by ART. Importantly, splenic organoids maintain excellent viability >85% for more than 14 days, allowing long-term experiments such as modeling ATI that were not possible in prior models. Upon ATI, robust virologic rebound is achieved within days, and transfer of allogeneic NK cells partially suppressed this virologic rebound. The ability to use secondary lymphoid organoids as a platform to test an NK cell-based cure strategy demonstrates a significant improvement over existing SLT explants.

Previous studies using SLT explants report the innate infectability of tissue CD4^+^ T cells compared to CD4^+^ T cells isolated from blood cells ^35–37,56^. Interestingly, the kinetics of infection differed between tonsil and spleen, with peak infection occurring 3 dpi in the tonsil and 7 dpi in the spleen. The continued increase in the frequency of infected cells in spleen organoids could be due to a variety of causes including (i) ongoing spread of infection to permissive cells, (ii) homeostatic proliferation of the infected cells (iii) a change in culture milieu that renders previously impermissive bystander cells to become HIV permissive, or (iv) cell-cell contact mediated infection ^17,43^. Notably, to achieve a similar level of infection in CD4^+^ T cells isolated from PBMCs, robust activation with anti-CD3, anti-CD28, and PHA along with infection with ∼10-fold more virus was required. This demonstrates the inherent differences in susceptibility of the CD4^+^ T cell population between blood and tissues.

Donor characteristics appear to influence the magnitude of HIV infection between organoids. We consistently observed higher levels of infection from the six pediatric tonsils tested compared to the adult tonsil and adult spleens. Future studies with tissues derived from age-matched donors will be needed to determine whether differing susceptibility to HIV infection is a result of the tissue inflammatory profile, age-dependent changes such as cellular senescence of CD4^+^ T cells that could limit the frequency of permissive cells, or a combination of these and other factors.

Both tonsil and splenic organoids recapitulated key features of natural infection, including a marked depletion of CD4^+^ T cells and infection of CM, EM, and T_fh_. This infection pattern is consistent with previous findings in PLWH ^11,15,16,40^. Unlike the HLAC explants ^39^, p24^+^ myeloid cells were also observed in both tonsil and splenic organoids. It is unclear whether these p24^+^ myeloid cells result from productive infection or the phagocytosis of previously infected T cells. Splenic organoids also consistently maintained viability above 85%, making splenic organoids a better model system for longer-term experiments.

As expected, targeting the late stage of the viral lifecycle with a protease inhibitor arrested CD4^+^ T cell depletion ^9,57^, reduced the frequency of p24^+^ cells, and suppressed the production of infectious virions. ART also replicated the post-treatment reduction in the inflammatory marker CXCL10 observed in PLWH ^47–49^. Although it takes several weeks to months to fully suppress HIV *in vivo* ^58^, in most donors, complete suppression of IP was achieved after 7 days of treatment, despite continued Gag p24 detection, consistent with low-level proviral expression in the absence of productive viral spread. This shortened timespan is highly advantageous for the rapid iteration for experimental timelines.

An additional advantage of the immune organoid model is the ability to visualize infected cells. When grown directly in chambered slides without an extracellular matrix, they adhered adequately and maintained their structure for fluorescence in situ hybridization. At 3dpi, we detected colocalization of vDNA^+^ cells actively transcribing vRNA. We also observed several vDNA^+^ cells that were not transcribing vRNA. These cells may represent early latently infected cells with replication-competent or defective proviral DNA ^6,50,57^. These imaging modalities will be powerful for visualizing the clearance of cells harboring vDNA and for testing novel cure strategies.

While PLWH experience rebound upon ATI, most *in vitro* studies induce rebound by mitogen stimulation. Here, viral rebound occurred when donor-matched uninfected splenocytes were added at ATI to mimic the migration of uninfected, HIV-susceptible cells to the site of infection, as would occur *in vivo* ^17,43^. We observed production of new viral particles within 3 days of ATI, a slightly shorter time scale than that observed in time-to-rebound studies conducted with PLWH, where rebound occurred within 5 days to 3 weeks ^7–10,12,57^. The difference in time to rebound between our *in vitro* system and *in vivo* studies could be due to several factors, including (i) the intracellular ARV concentration ^12^, (ii) time to ART initiation ^9,10^, (iii) individual participant reservoir size ^7–10,12,57^, or (iv) insufficient establishment of a reservoir during the seven day infection time. However, the detection of several hundred cells harboring intact viral genomes in cultures treated with ARV at the time of infection and maintained for 3 weeks, indicates that reservoir cells were generated. We demonstrated that in our model rebound can occur simply in the presence of new target cells, without the need for exogenous mitogens or stimulation. Given that our data shows a only small subset of CD4^+^ T cells are susceptible to infection and the recrudescence of IP in the presence of uninfected target cells, this could provide an alternative hypothesis as to why some people, also known as elite controllers, seem to control infect without the need for ART^59^. Perhaps in addition to their enhanced immune response to HIV infection^59^, they also do not produce the cells susceptible to HIV at high enough frequency to continually propagate the infection after the initial burst of viremia, allowing them to maintain a low or undetectable viral load. Our immune organoid model will allow further investigation to identify the factors that make tissue resident cells susceptible to productive infection and if the depletion of that cell population can act as a functional cure, like that of elite controllers.

Finally, immune organoids provide a platform for evaluating cure strategies. NK cells are cytolytic lymphocytes with an innate ability to kill virally infected and tumor-transformed cells through activating receptors on their cell surface and by engaging with activating ligands on target cells ^21,60^. We recently showed that NK cells can target and kill HIV-infected cells in a 2-D cell culture system via NKG2D and NKp30 ^21^. Prior NHP and human studies have correlated the presence of CXCR5^+^ NK cells with a decrease in HIV vDNA^+^ cells in lymph nodes, further implicating an active role of NK cells in HIV control in blood and tissues ^61,62^. These findings, in conjunction with a recent humanized mouse study that found that the transfer of allogenic NK cells delayed viral rebound and reduced the number of viral clones ^23^. IL-15 is an essential cytokine for NK cell survival; however, it is also a potent mitogen that can stimulate CD4^+^ T cells and induce viral replication, as observed here and by others ^63^. IL-15 increased the intact reservoir size by 2.6-fold relative to the infected vehicle-treated group. Despite this expansion of infected cells, weekly transfer of allogeneic NK cells reduced the number and frequency of p24^+^ CD3^+^CD8^-^ T cells, the number of intact viral genomes, and the production of IP in most organoid donors. This provides further evidence supporting the development of allogeneic NK cell-based cure therapies to target and eliminate the viral reservoir. However, if allogeneic NK cell-based strategies are pursued, they will need to circumvent the requirement for exogenous IL-15 by developing NK cell therapeutics that are longer-lived and do not require IL-15.

Overall, this human immune organoid platform provides a manipulatable, highly reproducible platform for studying HIV infection, treatment interventions, and cell-based therapies, thereby significantly benefiting both basic and translational research. This organoid model, initially developed to evaluate influenza vaccine responses, could also be used to assess HIV vaccine candidates, hopefully shedding light on why some vaccine candidates that looked promising in NHP models had less success in human trials ^64^. Overall, this platform can be broadened to evaluate the efficacy of next-generation ART and LRA function in tissues, and cytotoxic cell-based cure strategies for PLWH.

## Methods

### Informed consent and tissue sample collection and pre-processing

Whole tonsils were obtained from adults and children undergoing tonsillectomy for obstructive spleen apnea or hypertrophy at Stanford Hospital. All samples were collected in accordance with the Stanford University Institutional Review Board (Stanford IRB-60741 and IRB-30837). All participants or legal representatives signed written informed consent.

Spleen samples were collected from deceased organ donors through Donor Network West (DNWest), an Organ Procurement Organization serving Northern California and Northern Nevada. DNWest identifies potential organ and tissue donors and secures authorization for the procurement of organs and tissues for transplantation and research. After recovery, spleen samples were immersed in the University of Wisconsin (UW) preservation media and stored on wet ice until processing. The projects’s procurement of donor samples was reviewed and approved by DNWest’s *<u>Ethics and Mission Committee</u>* and endorsed by the Medical Advisory Board.

Tonsil samples were collected immediately after surgery and incubated in Ham’s 12 medium (Gibco, #31-765-092) containing Normomicin (Invivogen, #NC9273499), Penicillin/Streptomycin/Amphotericin B (#15240062) for at least 30 minutes at 4°C before processing. Afterwards, tonsils were manually sectioned into small pieces (∼5mm x 5mm x 5mm) and passed through a 100 mM filter using a syringe plunger to obtain a cell suspension. Filter was repeatedly washed with complete immune organoid medium (OM1) (RPMI with glutamax, 10% FBS, 1X Penicillin/Streptomycin, 1X Normomicin, 1% Insulin/transferrin/selenium supplement (Gibco, #41400045), 1X non-essential amino acids, 1X sodium pyruvate) to remove cells. Debris was reduced by Ficoll density gradient separation. Collected cells were washed with PBS, enumerated, and frozen into cryovials in FBS + 10% DMSO. Frozen aliquots were stored at −180°C until use.

Spleen samples were incubated in Ham’s 12 medium containing Normomicin and Penicillin/Streptomycin for at least 30 minutes at 4°C before processing. Spleen tissue was manually sectioned into small pieces and incubated in RPMI with glutamax, 10% FBS, 1X Penicillin/Streptomycin, and 1mL Normocin together with digestion enzymes (Collagenase IV [Worthington, #LS004188] and DNase I [Sigma, #04536282002]) and dissociated using the GentleMACS tissue dissociator (Miltneyi Biotec). After dissociation, spleen tissue was passed through a 100 mM filter using a syringe plunger to obtain a cell suspension. Filter was repeatedly washed with PBS containing FBS and EDTA. Red blood cells (RBCs) were lysed with ACK lysis buffer (KD Medical, RGF-015). Remaining RBC and granulocytes were depleted by magnetic isolation using EasySep Direct Human PBMC isolation kit (StemCell, #19654). Collected cells were washed with OM1, enumerated and frozen into cryovials in FBS + 10% DMSO. Frozen aliquots were stored at −180°C until use.

Healthy, deidentified donor leukoreduction system (LRS) chambers were purchased from Stanford Blood Center. Because these are fully de-identified, the Stanford IRB has determined that they do not constitute human subjects research. Peripheral blood mononuclear cells (PBMCs) were isolated from the LRS chambers by density gradient centrifugation using Ficoll-Paque PLUS (GE Healthcare), and cryopreserved in 10% DMSO (Sigma Aldrich) and 90% FBS (Thermo Fisher Scientific).

### Organoid preparation and infection

Organoid development has been previously described ^45^. Briefly, tonsil and spleen single cell suspensions were thawed in OM1 media supplemented with 1μl of benzonase per 10ml of OM1. Cells were pelleted at 1400rpm for 7 minutes at 4°C and resuspended in OM1 supplemented with benzonase. Subsequently, they were incubated for 30 minutes to 1 hour at 37°C in an incubator with 5% CO2. After incubation, cells were pelleted and resuspended in PBS for counting. After counting, the cells were pelleted a final time and resuspended in OM1 at a concentration of 6×10^6^ cells per 100 μL. Before cells were plated in Transwell inserts, 1 mL of OM2 medium (OM1 supplemented with 0.5 μg/ml recombinant human B cell-activating factor (BAFF) (BioLegend) was aliquoted into a standard 12-well plate. 100 µL of cell suspension was plated in 0.4-µm pore size PTFE transwell (EMD Millipore, # PICM01250) or in a 4-well Nunc Lab-Tek II Chamber Slide (Thermo Fisher Scientific, #154526PK).

For organoid infections, the multiplicity of infection (MOI) was based on the average frequency of CD19^-^ cells in the tissues, 20% for tonsils and 40% for splenic organoids, based on the frequency of CD19- cells (Supplementary Fig. 1) and previous work ^45^. After plating, appropriate MOI of Q23 virus was added directly onto the cells in the transwells. The 10 mM Tipranavir solution in DMSO (MedChemExpress, # HY-15148) and DMSO (ThernoFisher Scientific) was used as a vehicle control. Tipranavir and DMSO were diluted to equal volumetric concentrations in OM2. For ATI conditions, DMSO was diluted in OM2 to equal the volumetric concentration of Tipranavir. IL-15 was diluted to a final concentration of 50 ng/mL in OM2 with DMSO. The culture medium OM2 in the 12-well plate was replaced every 3-4 days with 1 mL of OM2 prepared with fresh BAFF.

For experiments in which donor-matched uninfected splenocytes were added, cells were harvested from the transwell by washing the cells off the transwell 2-5 times with 200 μL of PBS, followed by an additional 1 mL of PBS. This cell suspension was then pelleted at 500 g for 5 minutes at 4°C. The PBS was discarded, and cell pellets were resuspended in 100 μL OM1. 50 µL was returned to the original transwells, and the remaining cell suspension was used for flow cytometry staining and/or frozen for CS-IPDA. Fresh vial(s) of donor match splenocytes were thawed as stated above and resuspended at 6×10^6^ cells/100 µL, 50 µl (3*10^6^) of this cell suspension was added to the appropriate transwells. For conditions that received NK cells, isolated NK cells cultured overnight were pelleted at 500 g for 5 minutes at 4°C to remove culture media. Cell pellets were resuspended at 1×10^6^ cells/100 µL in OM1, and 50uL (5*10^5^) of NK cell suspension was added directly into the transwell.

### Cell lines

293T cells purchased from American Type Culture Collection (ATCC) (ATCC, #CRL-3216) were cultured in D10 (DMEM media supplemented with 10% FBS (Thermo Fisher Scientific) and 1% (PSA) penicillin/streptomycin/amphotericin (Thermo Fisher Scientific). TZM-bl cells were obtained from BEI Resources, NIAID, NIH (BEI Resources, #HRP-8129) and cultured in D10.

### HIV preparation and titer

All experiments were conducted using a full-length, replication-competent HIV-1 subtype A strain, Q23-17 (Q23), obtained from BEI Resources. This strain was isolated from a Kenyan woman approximately 270 days after estimated HIV acquisition ^46^. 293T cells were transfected with Fugene 6 (Promega) and full-length Q23 plasmid (BEI Resources, ARP-12649) in T225cm flasks. 293T supernatant was harvested 48 hours post-transfection, aliquoted on top of 20% sucrose gradient, and ultracentrifuged at 25,000g for 2 hours at 4°C. After ultracentrifugation, viral pellets were resuspended in PBS, aliquoted, and stored at −80°C. Viral stocks were titrated on TZM-bl cells (BEI Resources, ARP-8129) cultured in D10 supplemented with 10ug/ml DEAE-Dextran (Sigma-Aldrich, #93556-1G). 48 hours post-infection, TZM-bl cells were fixed with 500 µl of fixing solution, 1% paraformaldehyde (Electron Microscopy Sciences) and 0.2% glutaraldehyde (Thermo Fisher Scientific) in PBS for 5 minutes at room temperature. The wells were washed twice with PBS. Cells were then stained with 300 µl of staining solution, final concentration of 0.004 M potassium ferrocyanide, 0.004 M potassium ferricyanide, 0.002 M MgCl_2_, and 0.4 mg/ml X-gal in PBS to produce blue cells. Blue cells were then counted to determine the number of infectious particles. Titer value was used to calculate MOI for infections.

### CD4 T cell isolation and infection

CD4^+^ T cells were isolated and infected as previously described ^21^. CD4 T cells were isolated by negative selection using the CD4^+^ T cells isolation kit (Miltenyi Biotec, #130-096-533). Isolated CD4^+^ T cells were cultured overnight with plate-bound anti-CD3 (Thermo Fisher Scientific, #14-0037-82) in the presence of 1 µg/ml anti-CD28/anti-CD49d (BD Biosciences, #347690) and 3 µg/ml PHA-L (Thermo Fisher Scientific, #00-4977-93), incubated for 48 hours. Activated CD4+ T cells were infected with Q23 at MOIs 1, 5, and 10 using Viromag magnetofection (OZ Biosciences) for 24 hours.

### NK cell isolation

PBMCs were thawed, and NK cells were isolated by negative selection using NK-negative selection kits (Miltenyi Biotec, #130-092-657). NK cells were cultured overnight in R10 medium (RPMI 1640, with 10% FBS, 1% L-glutamine, 1% sodium pyruvate, and 1% PSA Penicillin/Streptomycin/ Actinomycin) supplemented with 300 IU/ml recombinant human IL-2 (R&D Systems) for 18hrs +/- 2hrs.

### Flow cytometry

Organoids were harvested from the transwells by rinsing the transwell with 200 μL of PBS 2-5 times. The harvested organoids were pelleted at 500 g for 5 minutes at 4°C, then transferred to a standard 96-well U-bottom plate for flow cytometry. Viability stain (ViaDye Red) (Table S1) and FC block (Biolegend, # 422302) was performed concurrently for 10 minutes at room temperature. 2X master mix of surface-staining antibodies (Table S1) was added to the cells in the presence of the viability and FC block mix for an additional 30 minutes. Cells were pelleted and washed twice in FACS buffer at 500g for 5 minutes at 4°C. Samples were fixed and permeabilized using the Foxp3 Transcription Factor Staining Buffer Set (Thermo Fisher Scientific, # 00-5523-00) according to the manufacturer’s instructions for 30 minutes to 1 hour. Intracellular staining for Ki-67 and Gag p24 (Table S1) was performed for 30 minutes at room temperature. Samples were collected on the Cytek Aurora spectral cytometer, and data were analyzed using FlowJo (v.10.10.0). Data visualization and statistical analysis were performed using RStudio (v. 4.3.2).

### TZM-bl Infectious viral particle quantification

The TZM-bl cell line was used to quantify HIV infectious particles via adaptation of previously published protocols utilizing this cell line to quantify HIV neutralizing antibodies ^65^. Supernatants from immune organoids were mixed with equal volumes of D10 and then serially diluted in a 1:1 mixture of OM2 and D10 to obtain 100 µl of each dilution in 96-well flat bottom plates. 10,000 TZM-bl cells suspended in 100 µl of D10 + 20 µg/mL DEAE-Dextran were then added to each well for a final volume of 200 µl/well and final concentration of 10 µg/mL DEAE-Dextran. After 44-50h of incubation at 37°C, the supernatants were removed and 50 µl of PBS was added to each well. 50 µl of britelite plus reagent (Revvity) was then added to each well, and after a 2-minute incubation, 90 µl from each well was transferred to the corresponding well of a black 96-well flat bottom plate with white backseal (Sigma-Aldrich, #Z732117) covering the bottom of the plate and read on a luminometer (Promega GloMax Discover). To interpolate the amount of virus in each sample, the luciferase values were compared to a seven-point standard curve of known quantities of HIV completed in technical triplicate and reduced using a five-parameter logistic (5-PL) equation in Prism (v. 10). For samples where multiple dilutions were able to be quantified, the most dilute concentration was taken and multiplied by the dilution factor to determine the concentration of HIV infectious particles for that sample.

### CS-IPDA

Intact and total HIV DNA in organoid samples was quantified via a cross-subtype intact proviral DNA assay (CS-IPDA), as previously described (Cassidy et al., 2022; Fish et al., 2022). Briefly, the CS-IPDA is a multiplex ddPCR that targets 3 regions of the HIV genome within the LTR/gag, pol and env genes. A reference RPP30/deltaD ddPCR run in parallel estimates cell counts and mechanical shearing, which occurs during gDNA extraction. CS-IPDA was performed in triplicate, and if no intact HIV proviruses were detected, additional replicates were performed until a minimum of 1e5 cells were interrogated. Results of the CS-IPDA and reference RPP30/deltaD assay were analyzed using RStudio with quality control measures as described ^66,67^.

### Simultaneous vRNA and DNA RNAscope-FISH

Organoids were fixed in 4% PFA for 30 minutes and washed 3 times in PBS for 5 minutes each. Organoids grown in transwells were sequentially embedded in 25%, 50%, 75% and 100% O.C.T compound (Electron Microscopy Sciences) for 20 minutes each. The tissues were frozen on dry ice, and additional OCT was added to the transwell to form a frozen puck. The frozen tissue puck was then dislodged from the transwell, placed in cryomold with additional OCT, and stored at - 80°C until sectioning. For organoids grown in the chambered slide, after fixation, cells were dehydrated by sequential immersion in 50%, 70%, 100% and 100% ethanol and stored at −80°C.

RNAscope was performed according to the manufacturer’s instructions with modifications. Samples were fixed in 10% neutral buffered formalin for 15 minutes at 4°C, then sequentially dehydrated in 50%, 70%, 100% and 100% ethanol for 5 minutes each. Tissues were air-dried, and hydrogen peroxide was applied to the sections for 10 minutes at room temperature, followed by washing twice with PBS for 2 minutes each. A hydrophobic barrier was drawn and allowed to air dry. Sections were treated with the manufacturer’s Protease III at a 1:15 dilution for 10 minutes at room temperature. Protease was removed, and slides were washed three times with PBS 2 minutes each. Tissues were immediately hybridized with either predesigned consensus Clade A sense probes to detect vDNA (ACD Bio-techne, #426341) or custom probes to detect vDNA (ACD Bio-techne, #1811141-C2, targeting reverse-complement sequence of 2541-4600) and vRNA (ACD Bio-techne, #1811131-C1, targeting 5791-8114 of Q23-17 genome). The custom vDNA-C2 probe was diluted 1:50 with the custom vRNA-C1 probe. Samples were hybridized with the probes in a humidified HybEZ oven (ACD Bio-techne) for 2 hours at 40°C. The probes were removed, and the slides were washed twice with 1X RNAscope wash buffer. Slides were stored in 5X saline sodium citrate (SCC) (Thermo Fisher Scientific) overnight. The following days, the slides were washed once with 1X RNAscope wash buffer, and the assay was continued. Probes were amplified sequentially per the manufacturer’s instructions, with AMP1 for 30 minutes, AMP2 for 30 minutes, and AMP3 for 15 minutes in a humidified HybEZ oven. Fluorescent signal was developed with RNAscope Multiplex FL v2 HRP-C1 for Clade A sense or custom vRNA-C1 probes, and RNAscope Multiplex FL v2 HRP-C2 was used for custom vDNA-C2 probe, along with TSA-vivid 520 diluted at 1:1500 in TSA buffer and TSA-vivid 650 diluted at 1:3000 in TSA buffer.

Immunofluorescence staining for CD4 was performed immediately after TSA signal development. Slides were washed three times with PBS for 5 minutes each. Tissues were blocked with blocking buffer containing PBS, with 0.1% triton X-100 (Thermo Scientific) and 1% BSA (Sigma-Aldrich) for 1 hour at room temperature in a humidified chamber. Blocking buffer was removed, and the slides were washed three times with PBS for 5 minutes each. Surface CD4 was detected using rabbit anti-CD4 (clone EPR6855, Abcam) conjugated to Alexa Fluor 555 or Alexa Fluor 647. CD4 antibody was diluted 1:200 in blocking buffer. Tissues were stained in a humidified chamber in the dark at 4°C overnight. The antibody was removed, and the slides were washed three times with PBS. Nuclei were stained with NucBlue Fixed Cell ReadyProbes Reagent (DAPI) (Thermo Fisher Scientific, #R37606) for 5 minutes at room temperature. Tissues were washed three times with PBS. Excess liquid was removed, and slides were mounted with Prolong Glass (Thermo Fisher Scientific, #P36980) with coverslip (Millipore Sigma, #CLS2980245). Images were captured on Leica Stellaris 8 DIVE upright Confocal multi-photon. Image analysis was performed in Fiji.

### Statistical analysis

Formal statistical tests were only performed when n ≥ 6. Unpaired T-test or Wilcoxon signed rank test were performed in the indicated experiments. Due to donor availability and labor constraints in the experiment, additional donors were not included for experiments when trends were apparent with n < 6. Data visualization and statistical analysis were performed using RStudio (v. 4.3.2).

## Acknowledgements

We would like to acknowledge the initial work of this project was funded by a pilot grant from the Stanford Innovative Medicine Accelerator (CAB). We would like to thank the participants and organ donors whose donations made the study possible, and the members of the Blish lab for critical input on the project like Radeesha Jayewickreme, Trisha Bernard, Leslie Chan, Izumi de los Rios Kobara, Xariana Vales Torres, and Soneida DeLine-Caballero.

## Author contributions

Conceptualization, S.A.S, J.E.S and C.A.B.; methodology, S.A.S, J.E.S, U.M.M, R.S.F and E.S; investigation, SAS, J.E.S, and R.S.F.; formal analysis, SAS, J.E.S, and R.S.F.; visualization, S.A.S; writing – original draft, S.A.S. and C.A.B; writing – review and editing, S.A.S, J.E.S, U.M.M, R.S.F, E.S, A.S, R.P, J.A.Z, J.T.K, D.A.L, M.M.D, C.A.B.; funding acquisition, D.A.L, J.A.Z, M.M.D, and C.A.B; resources, E.S, A.S, and M.M.D.; supervision, C.A.B.

