## Supplemental figures and table for "Modeling HIV infection, treatment, rebound, and intervention in human immune organoids"

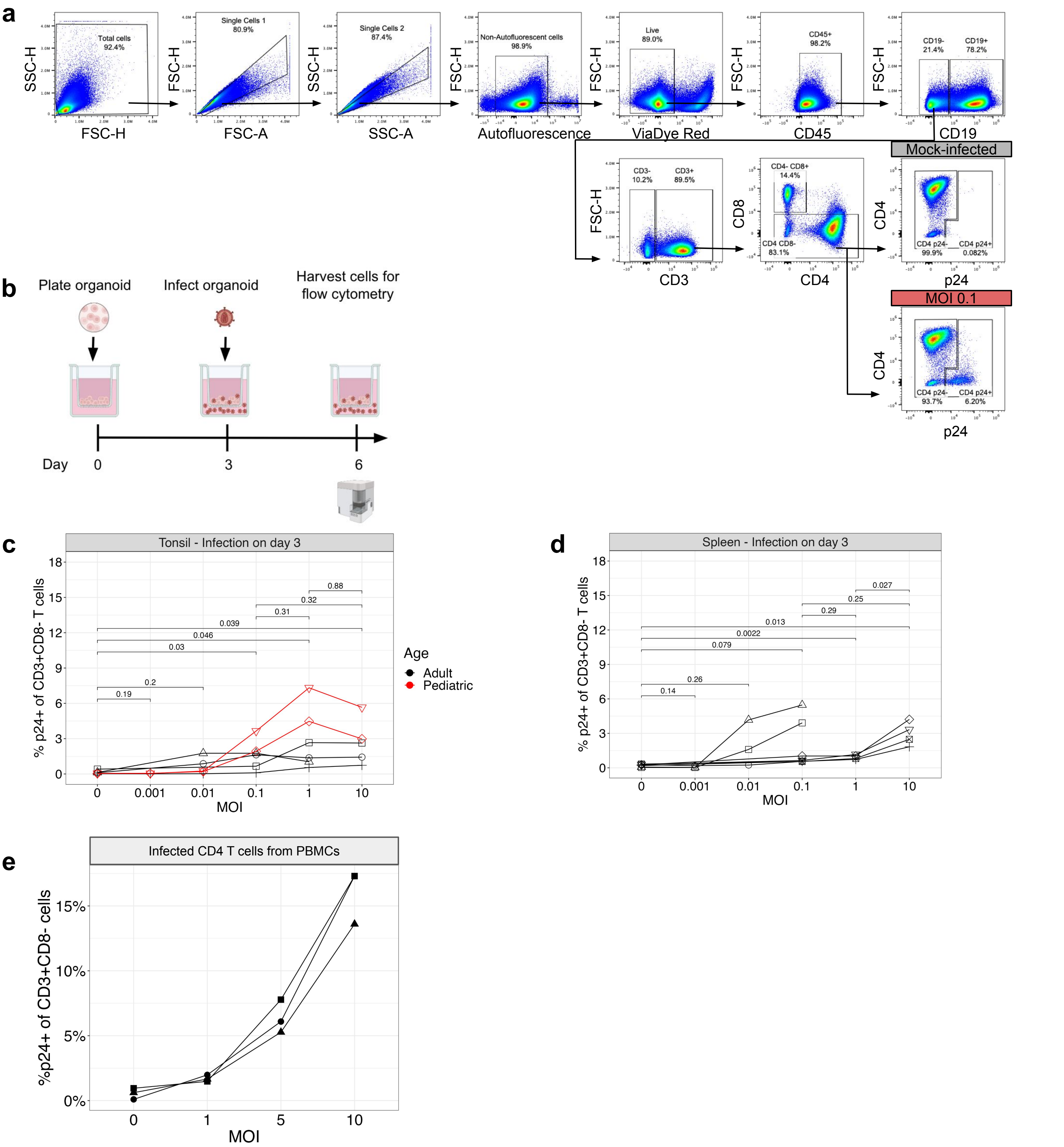


Supplemental Fig. 1: Limited population of tissue cells is permissive to HIV infection

**a**, Representative gating scheme for p24+ CD3+CD8- T cells. **b**, Schematic of infection timeline. Organoids were plated on day 0, infected on day 3, and then harvested for flow cytometry 3 dpi. **c-e**, Titration of MOI in, **c**, tonsil, **d**, splenic organoids and, **e**, PBMCs, measuring frequency of HIV Gag p24+ CD3+CD8- T cells at 3 dpi. Tonsil organoids were derived from 6 donors, 4 adult (black) and 2 pediatric (red). Splenic organoids were derived from 7 adult donors. CD4+ T cells isolated from PBMCs. Each symbol represents a donor. Statistical analysis were performed with unpaired t test.

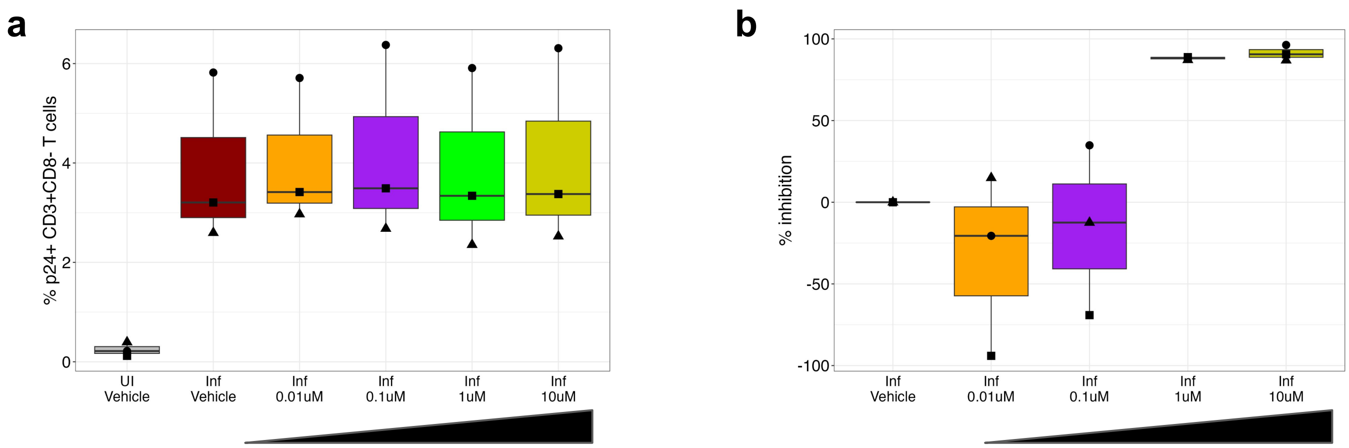


Supplemental Fig. 2: Tipranavir titration
Titration of tipranavir 0.1uM to 10uM on infected splenic organoids, 3 days post infect and treatment. Top row indicates infection status, bottom row indicates the TPV dose. **a**, Frequency of p24+ CD3+CD8- T cells. **b**, Quantification of inhibition of infectious viral particles in organoid supernatant. Each symbol represents a donor and is the mean of technical duplicates.

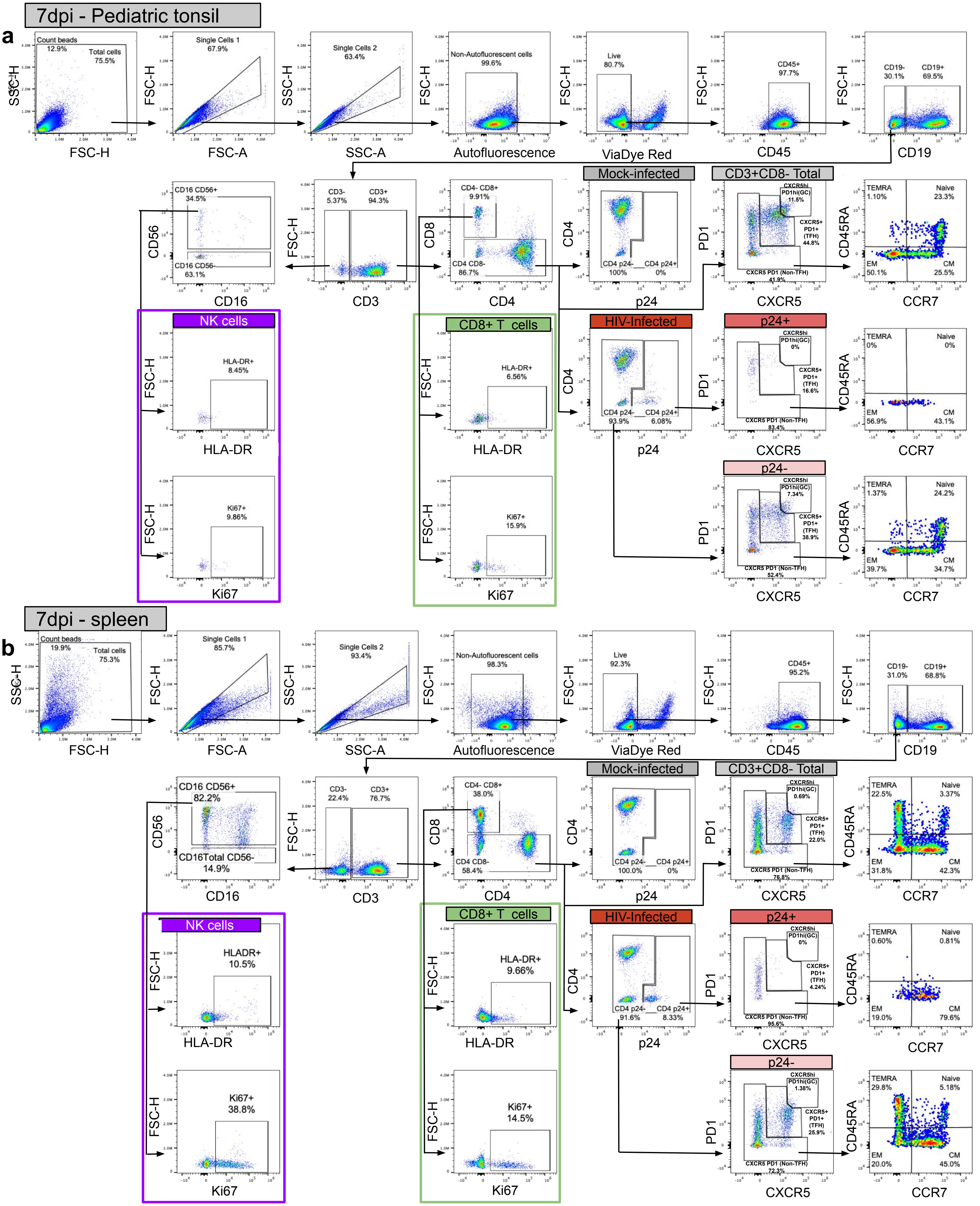


Supplemental Fig. 3: Representative gating scheme for, **a**, pediatric tonsil and, **b**, adult spleen, at 7dpi. Gating for p24+ CD3+CD8- T cell, CD3+CD8- T cell subsets, and CD8+ T and NK cell HLA-DR and Ki-67.



Supplemental Figure 3

### Supplemental Figure 4

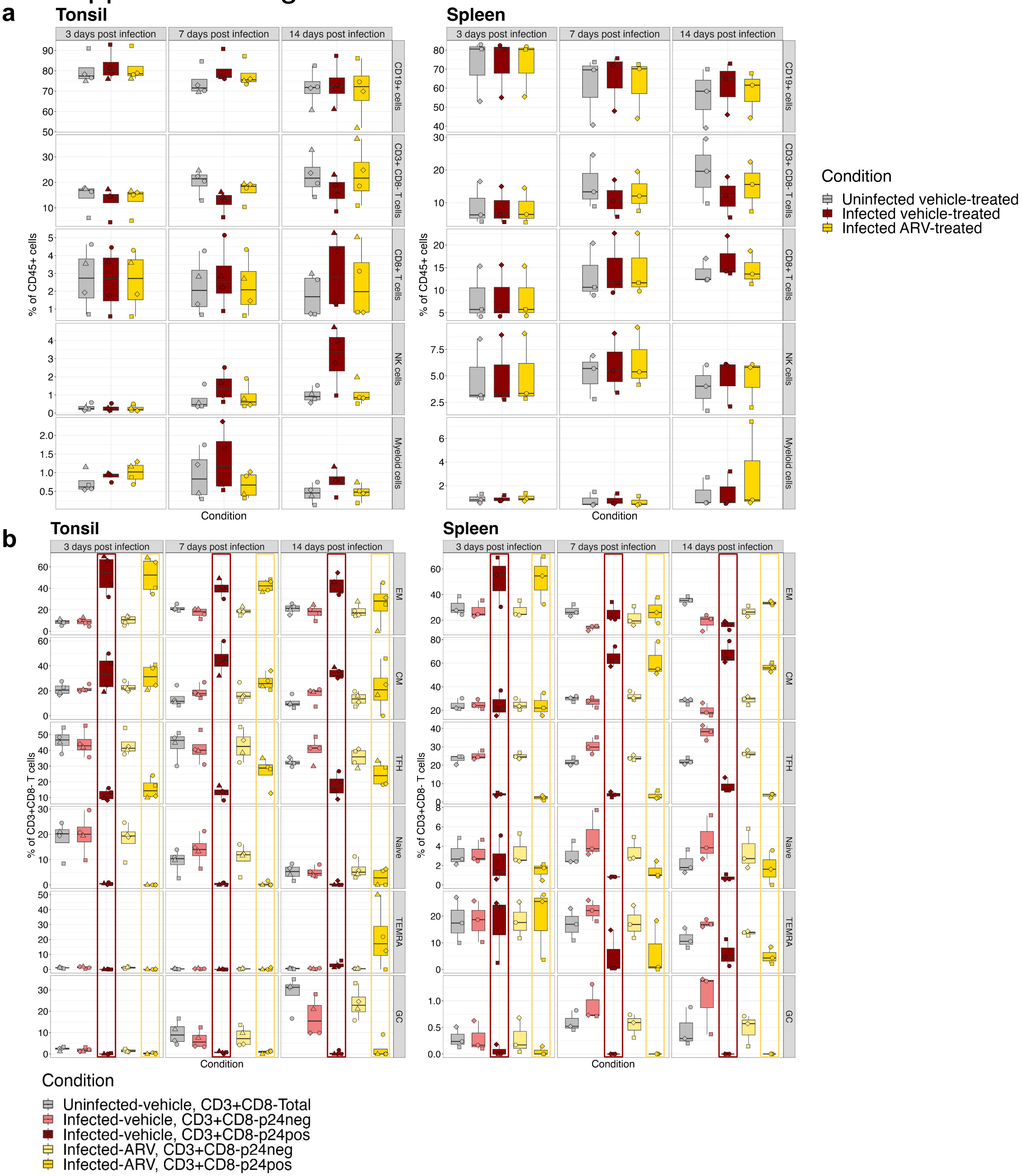


Supplemental Fig. 4: CM, EM and TFH are the major subsets of HIV p24+ CD3+CD8- T cells. 14 day time course infection of tonsil and splenic organoids. **a**, Frequency of the major immune cell populations CD19+ B cells, CD3+CD8- T cells, CD3+CD8+ T cells, NK cells and myeloid cells. **b**, Frequency of CD3+CD8- T cells subsets, stratified by uninfected CD3+CD8- T cells, infected vehicle-treated (p24- and p24+), and infected ARV-treated (p24- and p24+) organoids. Tonsil organoids were derived from 4 pediatric donors, splenic organoids were derived from 3 adult donors. Each symbol represents a donor and is the mean of technical duplicates.

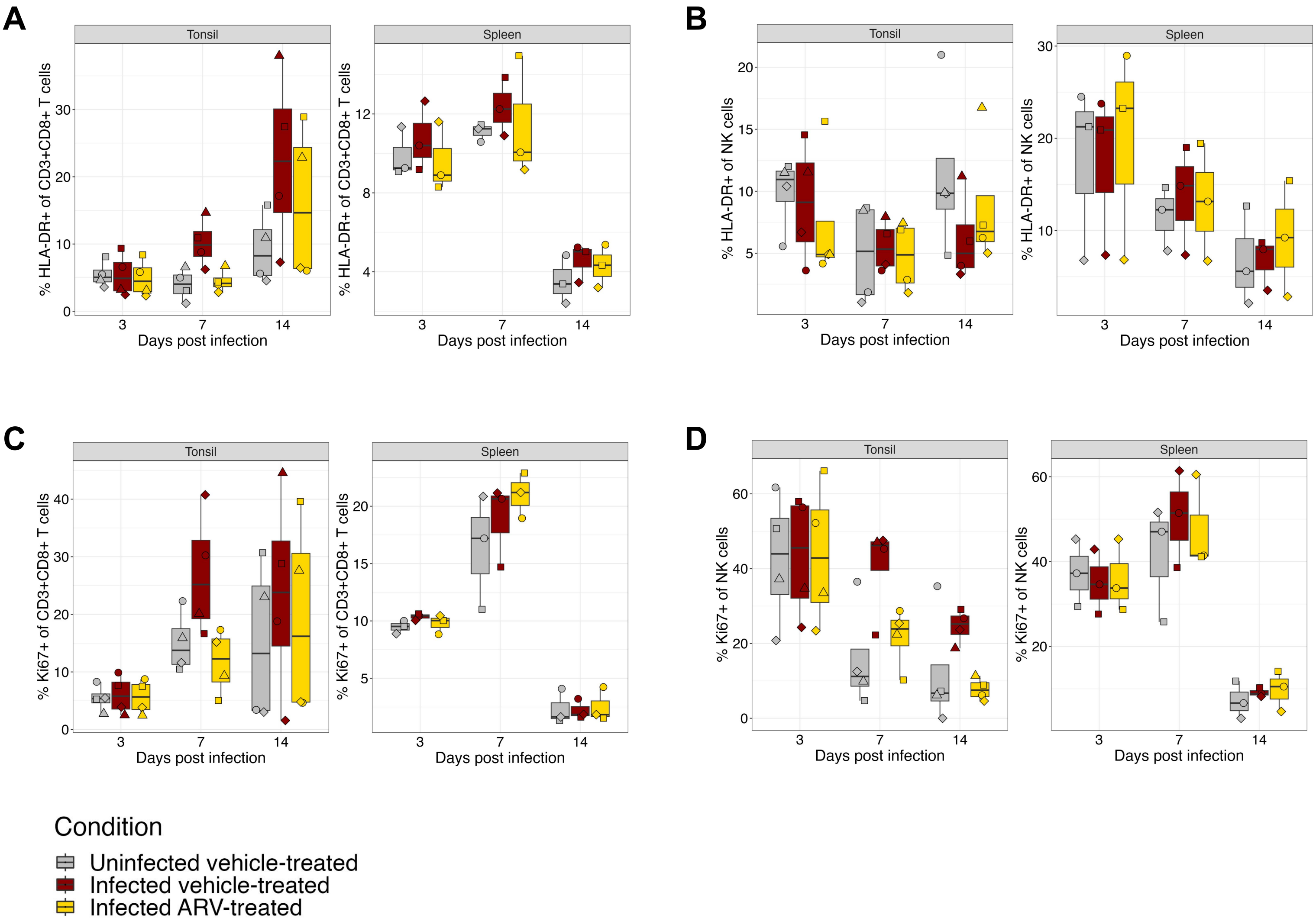


Supplemental Fig. S5: CD8+ T cells and NK cells are activated and proliferative in response to infection. Frequency of tonsil and splenic, **a**, HLA-DR+ of CD3+CD8+ T cells, **b**, HLA-DR+ of CD3+CD8+ NK cells, **c**, Ki67+ of CD3+CD8+ T cells, **d**, Ki67+ of CD3+CD8+ NK cells during 14 day time course. Tonsil organoids were derived from 4 pediatric donors, splenic organoids were derived from 3 adult donors. Each symbol represents a donor and is the mean of technical duplicates.

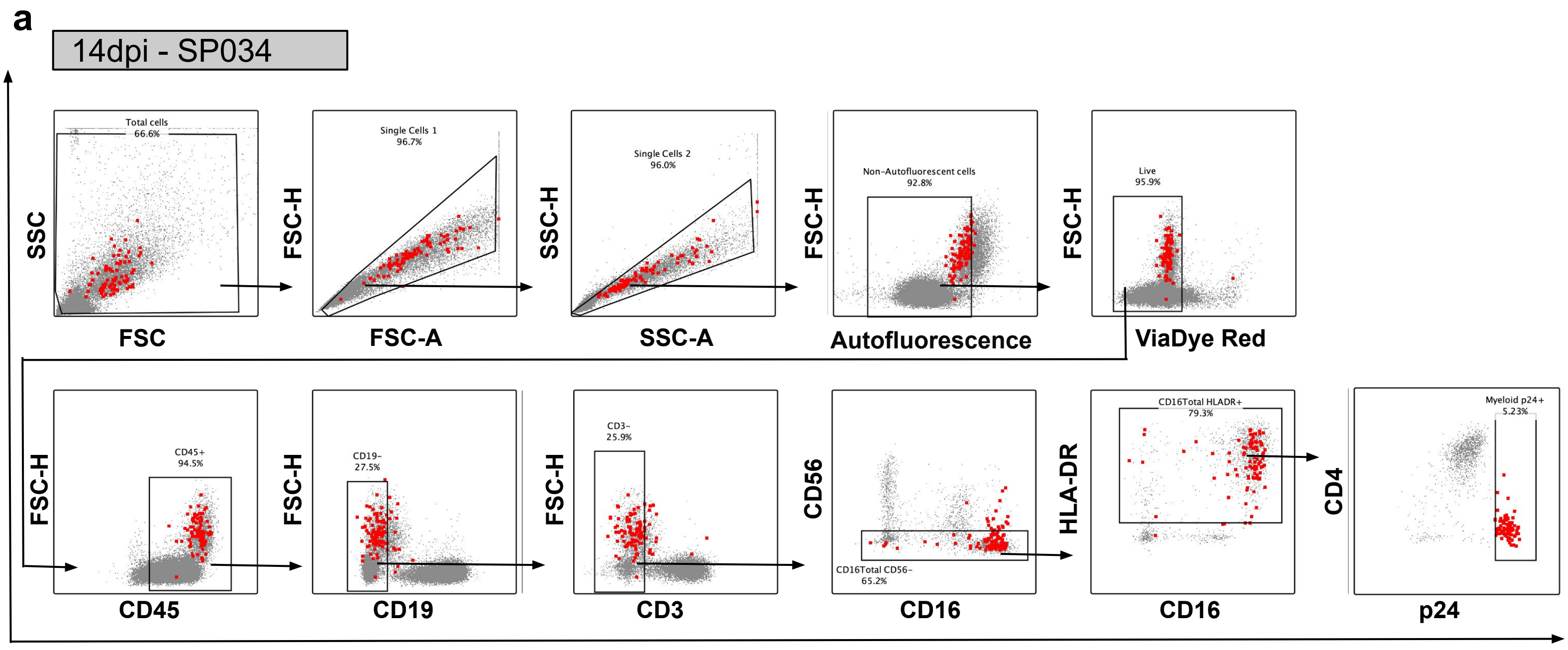


Supplemental Fig. 6:Gating scheme to identify p24+ myeloid cells in spleen.
**a**, Representative backgate plots for p24+ myeloid cells from infected, vehicle-treated culture at 14 dpi.

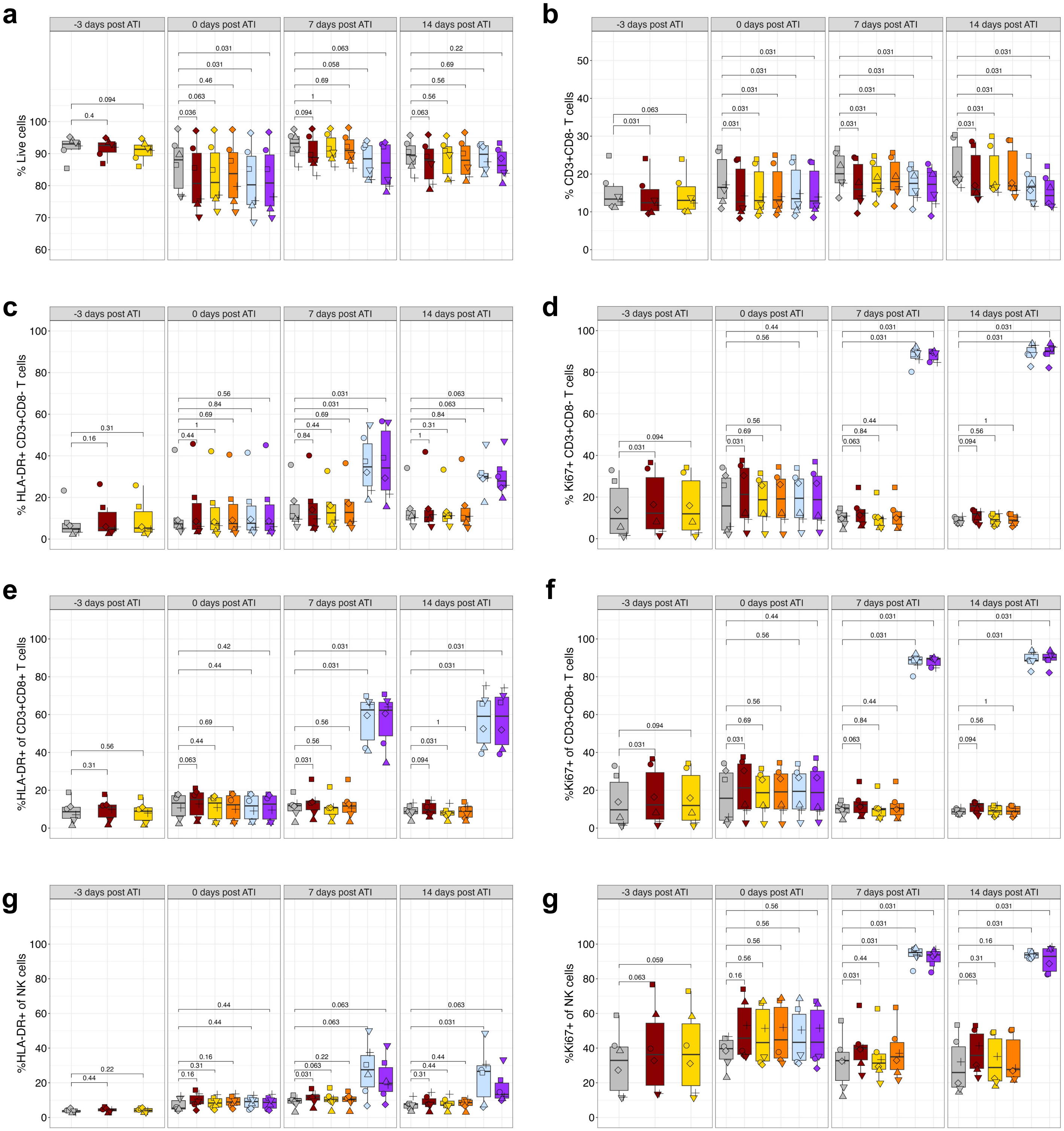


Supplemental Fig. 7: IL-15 treatment activates CD3+CD8- T, CD8+ T and NK cells.
ATI of HIV-infected splenic organoids followed by 14 days of allogeneic NK intervention. Frequency of, **a**, Live, **b**, CD3+CD8- T cell, **c**, HLA-DR+ of CD3+CD8- T cells, **d**, Ki67+ of CD3+CD8- T cells, **e**, HLA-DR+ of CD3+CD8+ T cells, **f**, Ki67+ of CD3+CD8- T cells, **g**, HLA-DR+ of CD3+CD8+ NK cells, **h**, Ki67+ of CD3+CD8+ NK cells during ATI with NK cell intervention. Splenic organoids were derived from 6 adult donors. Each symbol represents a donor and is the mean of technical duplicates.

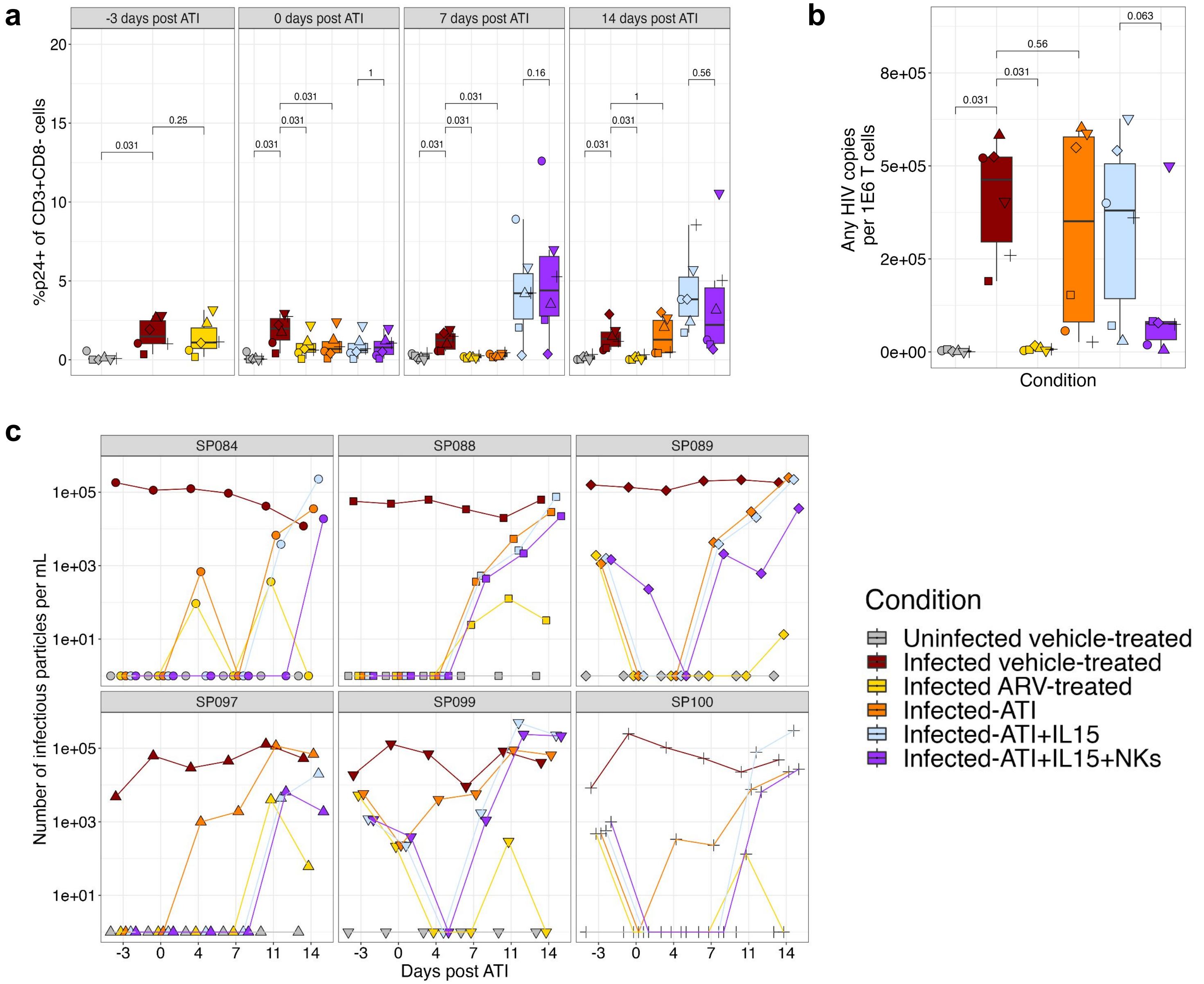


Supplemental Fig. 8: NK cell treatment reduces HIV reservoir and suppresses viral replication
ATI of HIV-infected splenic organoids followed by 14 days of allogeneic NK intervention. **a**, Frequency of p24+ CD3+CD8- T cells. **b**, Quantification of total copies of HIV by CS-IPDA. **c**, Quantification of the number of infectious viral particles in the supernatant of the organoids by individual donor during ATI, with and without NK cell intervention. Splenic organoids were derived from 6 adult donors. Each symbol represents a donor and is the mean of technical duplicates. Statistical analysis in were performed with wilcoxon signed rank test.

| Marker | Fluorophore | Working titer (uL/100uL) | Clone | Vendor |
|---|---|---|---|---|
| Viability | ViaDye-Red | 1:10,000 | N/A | Cytek |
| CD3 | BV510 | 2.5 | OKT3 | Biolegend |
| CD3 | BV650 | 1 | OKT3 | Biolegend |
| CD4 | BV421 | 1 | OKT4 | Biolegend |
| CD4 | PE/Dazzle 594 | 0.25 | SK3 | Biolegend |
| CD8 | cFluor V547 | 0.25 | SK1 | Cytek |
| CD8 | APC | 1.25 | SK1 | Biolegend |
| CD16 | BV480 | 0.5 | 3G8 | Biolegend |
| CD19 | PE | 1 | HIB19 | BD Biosciences |
| CD19 | cFlour BYG710 | 0.5 | HIB19 | Cytek |
| CD45 | Percp/Cy 5.5 | 1 | 2D1 | Biolegend |
| CD45 | cFluor B548 | 2 | 2D1 | Cytek |
| CD45RA | cFlour R685 | 5 | HI100 | Cytek |
| CD56 | APC | 0.25 | TULY56 | BD Biosciences |
| CCR7 | BV785 | 5 | G043H7 | Biolegend |
| CXCR5 | BV711 | 0.5 | J252D4 | Biolegend |
| HLA-DR | APC-H7 | 0.25 | G46-6 | BD Biosciences |
| PD-1 | PE-Cy7 | 0.5 | EH12.2H7 | Biolegend |
| Ki-67 | BV421 | 0.5 | ki-67 | Biolegend |
| p24 | FITC | 0.5 | HC-57 | Beckman Coulter |

Supplemental Table 1: Antibody resources for flow cytometry.
